# Gradient-Based Regulation of Activin A Directs Multilineage Liver Organoid Development from Human Pluripotent Stem Cells

**DOI:** 10.1101/2025.11.27.690974

**Authors:** João P. Cotovio, Alice C. Borges, Diogo E.S. Nogueira, Carlos A. V. Rodrigues, Joaquim M. S. Cabral, Tiago G. Fernandes

## Abstract

Morphogen gradients critically regulate the coordinated emergence of endodermal and mesodermal lineages during liver bud formation, yet current human hepatic organoid models inadequately replicate these spatial cues. Here we demonstrate that precise modulation of Activin A concentrations within a dynamic 3D vertical-wheel bioreactor system enables the co-emergence and spatial organization of hepatic epithelium and septum transversum mesenchyme-like populations from human pluripotent stem cells. High Activin levels induce uniform definitive endoderm and enhanced hepatic maturation, whereas low levels promote epithelial tube–like structures encased by mesenchymal cells, recapitulating early liver diverticulum architecture. This gradient-based approach autonomously generates parenchymal and non-parenchymal hepatic lineages, providing a scalable platform to model human liver organogenesis and engineer ventral foregut–derived tissues.

## Introduction

The development of liver organoids (LOs) offers tremendous potential for modeling liver biology, disease, and therapeutic responses^1–13^. However, current *in vitro* models remain limited in their ability to recapitulate the complex tissue interactions and structural organization seen during liver development. Most models focus primarily on generating liver parenchymal cells^14^ (the functional cells of the liver, such as hepatocytes) without adequately representing the dynamic interplay between endodermal and mesodermal tissues, which is critical for liver bud formation^15–17^.

An emerging strategy in organoid development involves the generation of liver organoids from a homogeneous population of human pluripotent stem cells (hPSCs) through a process known as co-emergence. In this approach, multiple cell types that constitute the developing liver bud arise and self-organize within a single hPSC-derived aggregate, recapitulating key aspects of *in vivo* liver development, where cell differentiation and spatial organization occur in a coordinated manner. Consequently, these liver organoids more accurately mirror the cellular diversity and structural complexity of the native developing liver.^18,19^.

Despite its promise, the full potential of the co-emergence strategy has yet to be fully realized. A major limitation lies in the need to optimize conditions that effectively drive the simultaneous differentiation and spatial arrangement of diverse cell lineages within a single organoid structure. One key biological principle guiding this process is the role of morphogen gradients in cell fate specification, where gene expression patterns are regulated in a concentration-dependent manner. Thus, a critical challenge for improving co-emergence lies in the precise modulation of signaling pathways and environmental parameters that influence morphogen diffusion and interpretation. Previous studies have shown that the activation of lineage-specific gene programs is tightly linked to the cellular distance from a morphogen source, as signal propagation occurs primarily through passive diffusion rather than active amplification mechanisms^20–23^.

In the context of liver development, the Transforming Growth Factor Beta (TGF-β) signaling pathway represents a compelling axis to explore, given its essential role in guiding endodermal differentiation^17,24^ . Building on this concept, we propose a novel strategy for the co-generation of septum transversum mesenchyme (STM) organoids, which provide a permissive microenvironment for the differentiation of definitive endoderm (DE) and hepatic epithelium^25^. It is well established that high concentrations of Activin A promote endodermal differentiation, while lower concentrations tend to induce mesodermal lineages^17^. To harness this concentration-dependent behavior, we differentiate hPSCs in a dynamic culture system characterized by low shear stress and controlled mass transport, enabling precise modulation of Activin A gradients to guide cell fate decisions^26–28^. Through this approach, we establish a refined co-emergence model that recapitulates key aspects of *in vivo* liver morphogenesis, allowing the synchronized differentiation and self-organization of multiple liver cell types from a single hPSC aggregate.

This strategy advances the field of organoid research by offering a more physiologically relevant model of liver development. Unlike traditional co-culture methods^29–32^, which are labor-intensive and often lack structural organization, the co-emergence approach enables spontaneous, coordinated formation of parenchymal and non-parenchymal cells, enhancing both reproducibility and scalability^33,34^. Additionally, this model holds broader implications for organogenesis studies and regenerative medicine, providing a robust platform for investigating developmental biology and improving disease modeling^34^. By leveraging fundamental principles of morphogen gradients and cell fate specification, particularly the role of Activin A in balancing endodermal and mesodermal differentiation, our approach represents a significant advancement in the generation of complex liver organoids. This work not only enhances our understanding of liver development but also lays the groundwork for more accurate models of organ function and pathology, with potential applications in drug discovery and personalized medicine.

## Materials & Methods

### Maintenance of hPSCs

Human induced pluripotent stem cells (iPS-DF6-9-9T.B, WiCell Bank, Wisconsin, USA) were maintained in mTeSR™1 medium (STEMCELL Technologies™) supplemented 1:200 (v/v) with penicillin/streptomycin (Gibco™, Thermo Fisher Scientific) on Matrigel-coated (Corning®) six-well culture plates in humidified incubators at 37°C and 5% CO_2_. Culture medium was changed daily. Cell passaging was performed every three to four days when reaching a confluency ∼70% using 0.5 mM EDTA (Invitrogen™, Thermo Fisher Scientific).

### Liver Organoid Differentiation

For liver organoid generation cells were seeded in a vertical-wheel bioreactor PBS Mini 0.1L (PBS Biotech,USA), 27 rpm, at a density of 250 000 cells/mL in 60 mL of mTeSR™1 supplemented with 10μM ROCKi Y-27632 (STEMCELL Technologies™). 24h after inoculation, full medium volume was replaced, and aggregates were maintained in mTeSR™1 without ROCKi at 25 rpm. For definitive endoderm differentiation, RPMI 1640 medium (Gibco™) supplemented with 1% (v/v) B-27™ minus insulin (Gibco™) and 1:200 (v/v) penicillin/streptomycin (Gibco™) was used as differentiation medium. On day 0 of differentiation, differentiation medium was supplemented with 10 ng/mL (A10) or 100 ng/mL (A100) of Activin A (PeproTech) and 6 μM of CHIR99021 (CHIR, Stemgent™) and agitation speed set to 30 rpm. After 24h, total media was replaced, and agitation speed changed to 33 rpm. For hepatic induction (day 3 of differentiation) cells were cultured in differentiation medium additionally supplemented with 10 ng/mL of FGF2 (PeproTech) and 20 ng/mL of BMP4 (PeproTech). From day 6 to day 20, cells were refreshed every three days with hepatocyte culture medium (HCM) BulletKit™ (Lonza) without the addition of human epidermal growth factor (hEGF). Medium was supplemented with 10 ng/mL oncostatin M (OSM, R&D Systems™), 0.1 μM dexamethasone (Dex, Sigma-Aldrich) and 20 ng/mL of hepatic growth factor (HGF, Sigma-Aldrich). For validation purposes, a 2D hepatic differentiation was performed adapting previously developed protocols from Takebe et al.^32^, Ang et al.^35^ and Loh et al^36^.

### Immunocytochemistry

Aggregates were fixed in 4% (m/v) PFA (Sigma-Aldrich®) at 4°C for 30 minutes. For cryosections, aggregates were then incubated in 15% (m/v) sucrose in PBS at 4°C overnight, embedded in a 7.5%/15% gelatin/sucrose mixture, and frozen in isopentane at -80°C. Samples were sectioned at 12 μm using a cryostat-microtome (Leica CM3050S, Leica Microsystems), collected on Superfrost™ Microscope Slides (Thermo Scientific), and stored at -20°C. Before staining, sections were de-gelatinized in PBS at 37°C for 45 minutes, incubated in 0.1 M glycine (Merck Millipore) for 10 minutes at RT to remove PFA residues and permeabilized with 0.1% (v/v) Triton X-100 (Sigma-Aldrich®) in PBS, 10 min, RT. Slides were blocked with 10% (v/v) fetal goat serum (FGS, Gibco™) in TBST (20 mM Tris-HCl pH 8.0, 150 mM NaCl, and 0.05% (v/v) Tween-20, Sigma-Aldrich®) at RT for 30 minutes and incubated with primary antibodies, 10% FBS/TBST at 4°C overnight. Following primary antibody incubation, secondary antibodies were added for 30min, RT, and nuclear counterstaining was performed using 1.5 μg/mL 4’,6-diamidino-2-phenylindole (DAPI, Sigma-Aldrich®) in PBS, 5min, RT. Images were acquired using a LSM 710 Confocal Laser Point-Scanning Microscope (Zeiss), and data analysis was conducted using ZEN Imaging Software (Zeiss) and ImageJ Software. List of primary antibodies and dilutions on Supplementary Table 1.

### Quantitative real time (qRT)-PCR

Total RNA from cells of dissociated liver organoids was extracted using High Pure RNA Isolation Kit following the provided instructions. RNA was quantified using a nanodrop and 1 μg of RNA was converted into cDNA with High Capacity cDNA Reverse Transcription Kit (Applied Biosystems™/ Thermo Fisher Scientific) also following the provided instructions. PCR reactions were run using SYBR Green Master Mix (Nzytech). Reactions were run in triplicate using ViiA™ 7 Real-Time PCR Systems (Applied Biosystems™/ Thermo Fisher Scientific) and data were analysed using QuantStudio™ Real Time PCR Software (Applied Biosystems™/ Thermo Fisher Scientific). The analysis was performed using the ΔΔCt method and values were normalized against the expression of the housekeeping gene glyceraldehyde-3-phosphate dehydrogenase (GAPDH). List of primers on Supplementary Table 2.

### RNA Sequencing

Total RNA from three different organoid differentiations was extracted at different differentiation stages using High Pure RNA Isolation Kit (Roche), according to manufacturer ’s instructions. RNA samples were processed and sequenced by NovoGene. Integrity was assessed using an Agilent 2100 Bioanalyzer followed by library preparation using NEBNext® Ultra TM Directional RNA Library Prep Kit for Illumina® and RNA sequencing using an Illumina NovaSeq 6000 system with a paired end 150 bp strategy. For RNA-seq analysis, sample run was aligned to hg38 reference genome using HISAT2 software. Counts were obtained using Featurecounts.

For validation, 2D differentiation libraries were prepared using Lexogen QuantSeq 3′mRNA-Seq Library Prep. Kit FWD for Illumina using standard protocols. Sequencing was performed using Illumina HiSeq (50 cycle’s protocol) or NextSeq (75 cycle’s protocol) platforms. Sample read quality, reads mapping and counting were performed by a standard protocol from BlueBee Genomics Platform (http://www.bluebee.com/). For each sample individually, the pipeline includes reads trimming using BBDuk, alignment against genomic sequence using STAR aligner, gene read counting using HTSeq and quality control using RSeQC.

### Data Analysis

DESeq2 R package was used to normalize raw count data and Differential Gene Expression analysis (DGE). Transcriptomic results present a p-value according to Wald statistical test and p-values were corrected using the Benjamin-Hochberg (BH) method. Genes were considered up or downregulated when a onefold or greater change was exhibited and whose expression values have statistically significance associated (p-value <0.05). Gene ontology (GO) and Gene Set Enrichment Analysis (GSEA) were both performed using the clusterProfiler R package. For GO analysis, p-value threshold was set to <0.05. GSEA used a p-value cut-off of 0.05 and p-values were corrected using the BH method. Deconvolution analysis was carried out using the DeconRNAseq package within the R environment. A reference signature matrix of pure fetal cell-type expression profiles was used (GSE156793_S6)^37^. Other reference data sets used include a fetal tissue data set (GSE156793_S4)^37^ and an adult liver data set^38^. Results from data analysis were visualized in plots created with the ggplot2 R package. In protein–protein interaction analysis differentially expressed genes between A10 and A100 conditions were analyzed using the STRING database (v11.5). Interactions were derived from functional and physical protein associations with a minimum required interaction score of 0.400 (medium confidence). Active interaction sources included text mining, experiments, databases, co-expression, neighborhood, gene fusion, and co-occurrence. The network was restricted to query proteins in the first shell only, and disconnected nodes were excluded. Visualization was performed in STRING, with edges weighted by confidence score and nodes clustered by functional pathway.

### Statistical Analysis

Statistical significance was determined using a mixed-effects analysis with GraphPad Prism9® for all quantification except differentially expressed RNAseq data. Data is represented as mean ± SD for at least three replicate samples (see figure legends for additional information).

### Data Accessibility

RNA-seq data for this study are available through Gene Expression Omnibus (GEO) Accession number GSE311252

## Results

### Induction of Hepatic Differentiation in a 3D Culture System

For differentiation into definitive endoderm (DE), TGF-β/Nodal/Activin signaling was activated by adding Activin A at either 10 ng/mL (A10) or 100 ng/mL (A100). This pathway plays a crucial role in driving cells through a mesendodermal stage, characteristic of the primitive streak (PS), and subsequently into definitive endoderm. WNT signaling was also activated using CHIR99021 (CHIR), which synergizes with TGF-β signaling to enhance differentiation efficiency.

Following DE induction, from day 3 to day 5, Fibroblast Growth Factor 2 (FGF2) and Bone Morphogenetic Protein 4 (BMP4) were added to promote hepatic specification. These factors mimic the inductive signals from the cardiac mesoderm and septum transversum mesenchyme (STM), guiding the endoderm towards early hepatic differentiation. From day 6 onwards, the medium was supplemented with Oncostatin M, dexamethasone, and Hepatic Growth Factor (HGF) to promote hepatocyte maturation and functionalization (Fig. 1A).

**Figure 1.**
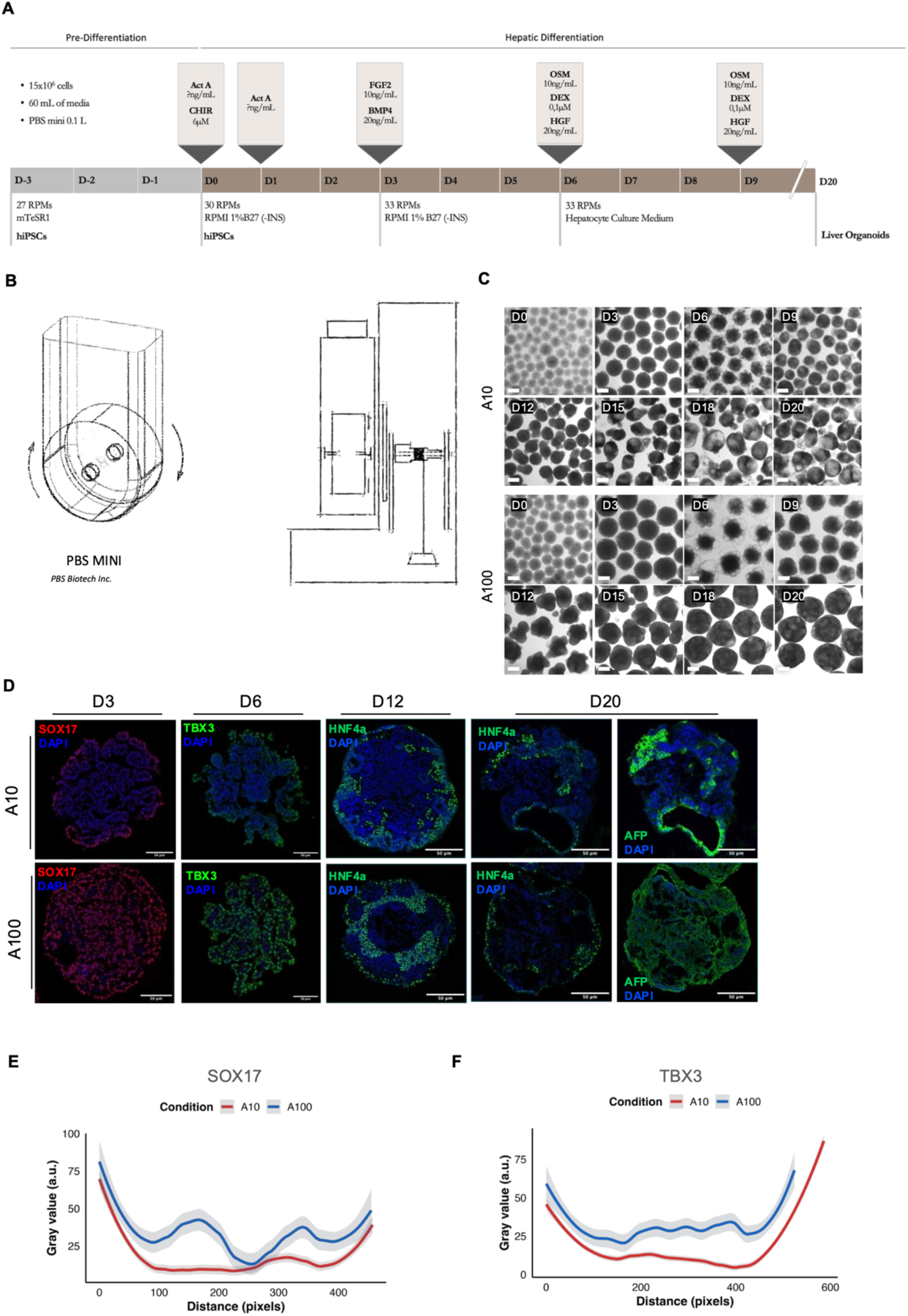
Phenotypic characterization of 3D hiPSCs-Derived Hepatic Organoids using a Vertical-Wheel Bioreactor (A) Schematic representation of 3D hepatic differentiation: Human induced pluripotent stem cells (hiPSCs) were directed through a three-step protocol: definitive endoderm (day 0–3), foregut/hepatic specification (day 3–6), and hepatic maturation (day 6–20). **(B) Schematic of the PBS Mini Vertical-Wheel bioreactor.** hiPSCs were seeded and differentiated in the PBS Mini Vertical-Wheel system (PBS Biotech Inc.). **(C) Aggregate morphology during hepatic differentiation:** Brightfield images show aggregate morphology under A10 and A100 culture conditions at selected time points from day 0 to day 20. Scale bars, 300 μm. **(D) Immunofluorescence of liver organoids at day 20.** Organoids differentiated under A10 and A100 culture conditions were stained for SOX17 (magenta), TBX3 (green), HNF4α (green), and AFP (green); nuclei were counterstained with DAPI (blue). Representative confocal images are shown, with top and bottom rows depicting independent organoids imaged under identical acquisition and display settings. Scale bars, 50 μm. **(E) Quantification of SOX17 staining intensity in liver organoids.** Staining intensity in organoids differentiated under A10 (red, n=1) and A100 (blue, n=1) culture conditions is shown. Curves represent LOESS-smoothed means with 95% confidence intervals; image brightness and contrast and pixel size were standardized before analysis. **(F) Quantification of TBX3 staining intensity in liver organoids.** Staining intensity in organoids differentiated under A10 (red, n=1) and A100 (blue, n=1) culture conditions is shown. Curves represent LOESS-smoothed means with 95% confidence intervals; image brightness and contrast and pixel size were standardized before analysis.

This phased protocol, which begins with TGF-β-mediated definitive endoderm induction followed by BMP and FGF signaling to promote foregut restriction and hepatic lineage commitment, is well established in the literature^14,39^.

To ensure uniform exposure to nutrients and signaling molecules, cells were seeded and maintained in a vertical-wheel bioreactor throughout differentiation (Fig. 1B). This platform supports homogeneous distribution of signaling molecules due to enhanced mass transfer and low shear conditions, ideal for the generation of uniform-sized aggregates ^27,40^ . Aggregate size was controlled to 280–300 μm in diameter, which is optimal for mesendoderm induction^26^. Aggregate morphology provided an early indication that different Activin A concentrations led to distinct outcomes. Aggregates treated with lower Activin A concentrations seem to develop increasingly prominent cystic regions, particularly noticeable during later differentiation stages. These structures were typically spherical or oval and likely lined by an epithelial cell layer. In contrast, aggregates treated with higher Activin A concentrations maintained a denser morphology throughout differentiation, culminating in an internal reticulated organization (Fig. 1C).

Drawing on previous 2D differentiation protocols (Supp. Fig. 1A), where expression of definitive endoderm markers *SOX17* and *FOXA2* peaked at days 3 and 7 respectively, we monitored the expression of *TBX3* (an early hepatic marker, D7–D9) and hepatoblast/hepatocyte markers (*HNF4α*, *CK19*, *AFP*, *ALB*) throughout the 3D differentiation stages to assess lineage progression under optimized conditions (Supp. Fig. 1B).

To evaluate whether these markers were also expressed in our 3D system, immunostaining was performed on cryosections at different time points. SOX17 was detected in D3 aggregates, TBX3 at D6, HNF4α from D12 onwards and AFP from D20 under both Activin conditions (Fig. 1D). These findings confirmed that our 3D culture recapitulates key hepatic differentiation milestones. Supporting the morphological observations from brightfield images, immunofluorescence (IF) revealed differences in spatial marker distribution across aggregates.

To determine whether early Activin A levels influenced downstream protein expression, we compared spatial staining profiles across aggregates. Under low Activin conditions, SOX17 and TBX3 showed pronounced peripheral enrichment (SOX17: 19.7 and 22.6 a.u.; TBX3: 19.3 and 13.0 a.u.) with reduced signal at the core (SOX17: 12.2 a.u.; TBX3: 9.9 a.u.). In contrast, high Activin exposure resulted in uniformly elevated expression throughout the aggregate (SOX17: 41.1, 25.1, and 27.0 a.u.; TBX3: 31.6, 27.4, and 37.1 a.u.), indicating a more homogeneous morphogen distribution (Fig. 1E–F).

### Activation of Regulatory Transcription Factors Highlights Network Specificity in Liver Organoid Development

To further validate the progression of hepatic differentiation, we investigated a transcription factor network previously described as essential for liver bud emergence. Gene Set Enrichment Analysis (GSEA) was performed on differentially expressed genes from day 3 (D3, definitive endoderm stage) and day 20 (D20, hepatic lineage) samples.

As expected, ontology terms related to TGF-β/Nodal/Activin and WNT (both canonical and non-canonical) pathways were enriched in both A10 and A100 conditions at D3 (Fig. 2A). The increased expression of classical TGF-β targets such as *SOX17* and *FOXA2* confirmed successful commitment toward endodermal fates (Fig 2C, Supp Fig 2A). Notably, *NODAL* expression differed between A10 and A100. In line with the higher activation dosage, A100 samples showed a marked peak of *NODAL* expression at D3, indicating a stronger induction of endodermal identity (Fig. 2B).

**Figure 2.**
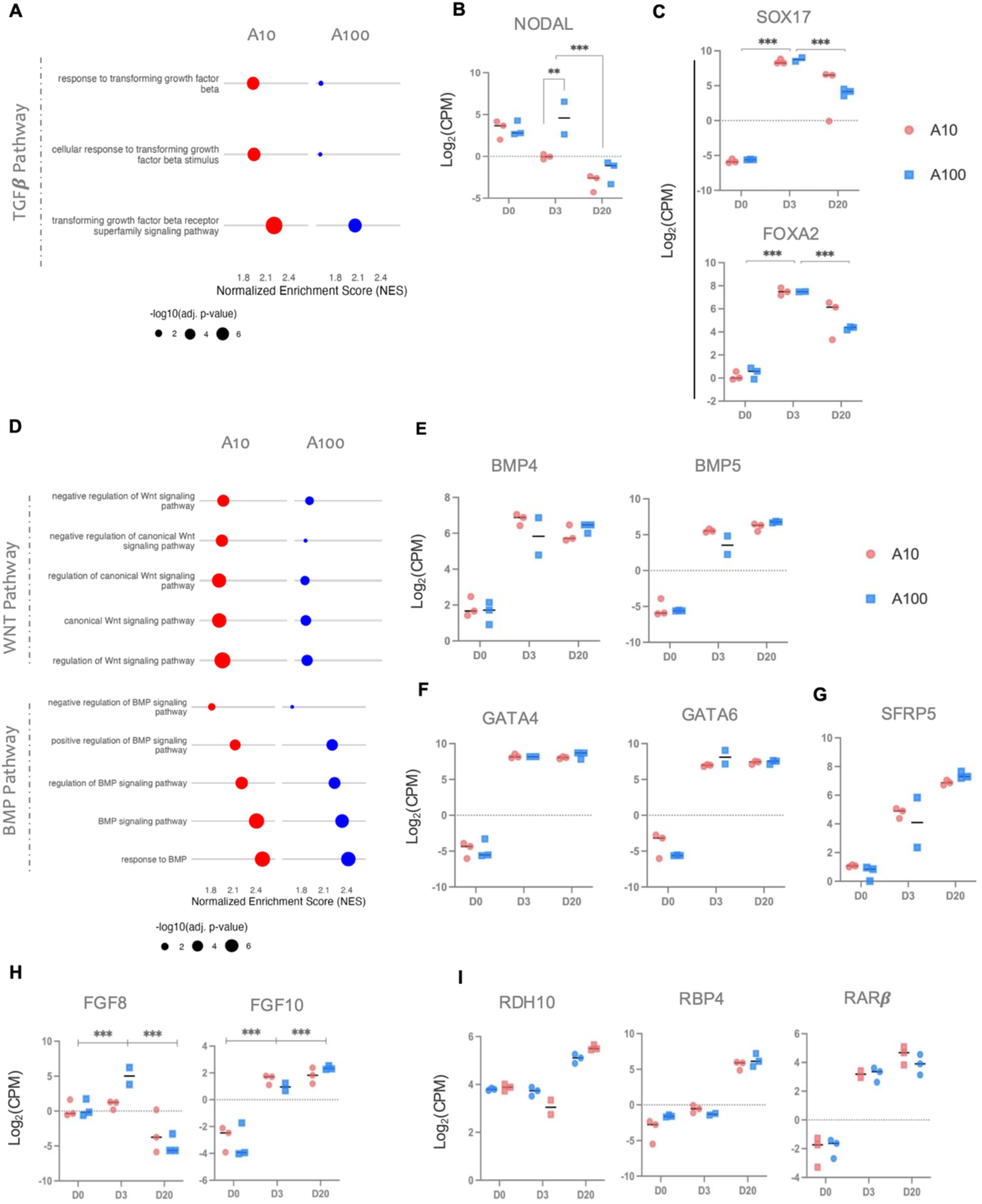
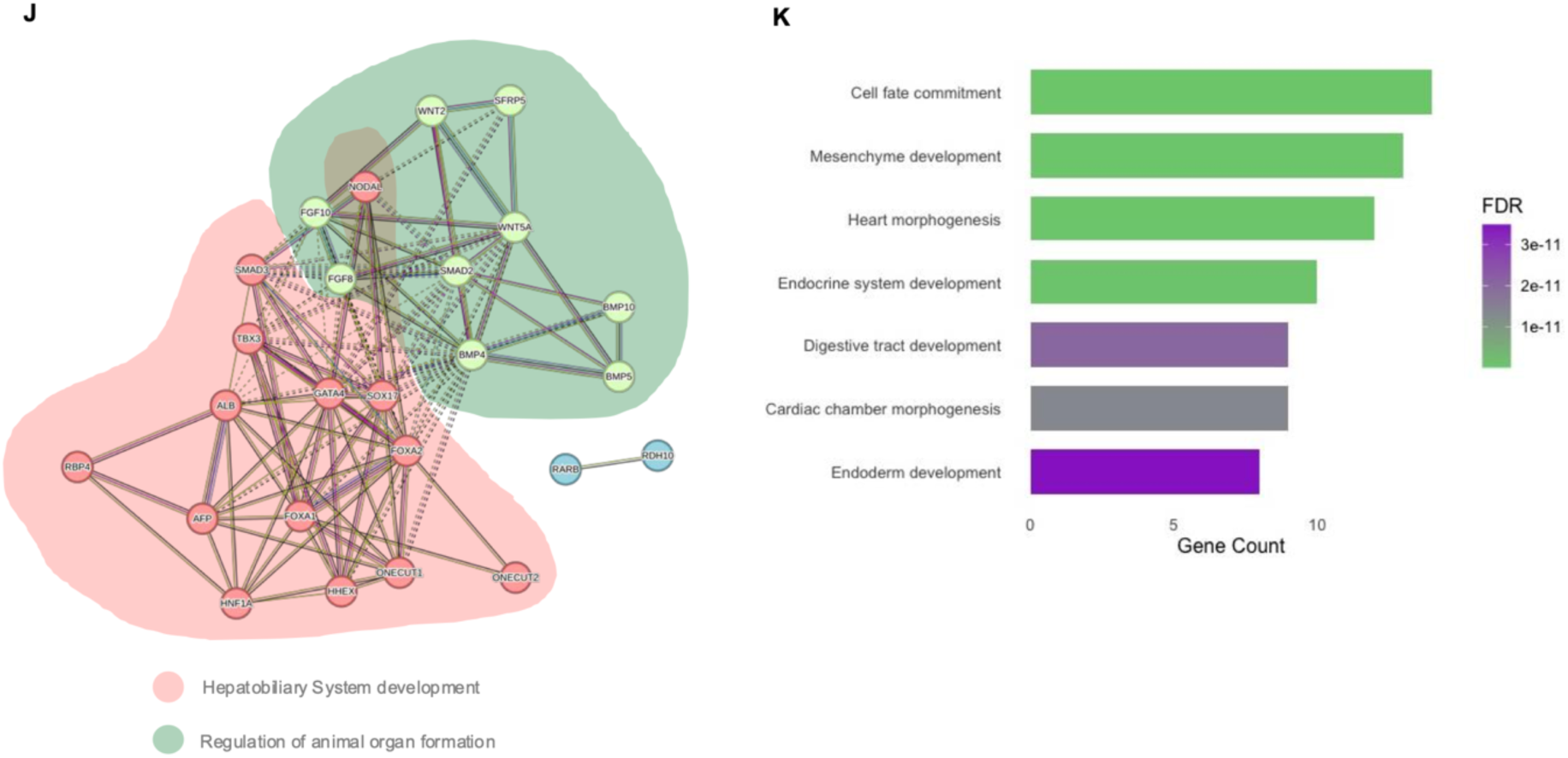
Integrated Pathway Activation and Network-Specific Transcription Factor Dynamics highlight regulatory drivers of liver organoid development in A10 vs A100 conditions. **(A) GSEA of TGF-β pathway–related GO terms between A10 and A100:** Dot plot of terms passing multiple-testing correction (adjusted *p* < 0.05). NES is plotted on the x-axis, GO terms on the y-axis, and dot size represents –log₁₀(adjusted *p* value). (B), (C) Expression dynamics of *NODAL*, *SOX17*, and *FOXA2* during differentiation: Normalized RNA-seq expression levels (log₂ counts per million, CPM) of *NODAL* (B), and *SOX17* and *FOXA2* (C) at day 0 (D0), day 3 (D3), and day 20 (D20) of differentiation. Circles represent A10 samples and squares represent A100 samples (biological replicates shown individually). Statistical significance was assessed using mixed-effects models. **P** values are indicated as follows: ** (P < 0.05), *** (P < 0.005). **(D) GSEA of WNT and BMP pathway–related GO terms between A10 and A100:** Dot plot of terms passing multiple-testing correction (adjusted *p* < 0.05). NES is plotted on the x-axis, GO terms on the y-axis, and dot size represents – log₁₀(adjusted *p* value). **(E – I) Expression dynamics of key pathway genes during differentiation:** Normalized RNA-seq expression levels (log₂ counts per million, CPM) of *BMP4* and *BMP5* (E), *GATA4* and *GATA6* (F), *SFRP5* (G), *FGF8* and *FGF10* (H) and *RDH10, RBP4* and *RARβ* (I) at day 0 (D0), day 3 (D3), and day 20 (D20) of differentiation. Circles represent A10 samples and squares represent A100 samples (biological replicates shown individually). Statistical significance was assessed using mixed-effects models. **P** values are indicated as follows: ** (P < 0.05), *** (P < 0.005). **(J) Protein–protein interaction network of differentially expressed genes:** Protein–protein interaction network constructed from differentially expressed genes between A10 and A100 conditions using the STRING database. Nodes represent proteins and edges indicate predicted functional associations, with edge thickness reflecting interaction confidence scores. Distinct colors denote functional clusters or pathways identified within the network. **(K) Gene set ontology enrichment of network-derived genes:** Gene Ontology (GO) enrichment analysis of differentially expressed genes from the protein–protein interaction network. Bars indicate significantly enriched biological processes and molecular functions (FDR < 0.05), with length corresponding to gene count.

Regulation of the BMP pathway, critical for hepatic induction, was also evident in both datasets. Expression of BMP ligands (Fig2E) and downstream effectors, including *GATA4* and *GATA6*, followed a consistent pattern, upregulated at D3 and sustained through D20 (Fig. 2F). Interestingly, despite differences in morphogen exposure, both conditions showed not only pathway activation but also enrichment for negative regulation terms related to WNT and BMP signaling (Fig2D). This is consistent with embryonic anterior–posterior patterning, in which gradients of BMP and WNT activity across the septum transversum mesenchyme and cardiac mesoderm are required to maintain foregut and hindgut boundaries^41^. Supporting this patterning process, we observed an increase in *SFRP5* expression from D3 onwards, with SFRP5 being a secreted WNT antagonist involved in foregut identity establishment^42,43^ (Fig2G). These findings suggest that negative regulation of WNT signaling is endogenously activated within our system, reinforcing anterior endoderm patterning.

Mesoderm-derived FGF ligands contribute to this spatial and temporal organization by further refining endodermal identity and promoting hepatic specification^43,44^. Expression analysis revealed that *FGF8* and *FGF10* were upregulated at D3, potentially representing mesodermal paracrine signals. Notably, *FGF8* expression declined by D20, consistent with a transient role in early liver development, while *FGF10* expression was maintained throughout, aligning with its known role in hepatoblast proliferation and expansion^44^ (Fig2H).

We also examined the involvement of retinoic acid (RA) signaling, an important mesenchymal cue during liver bud expansion. Despite the absence of exogenous RA in our protocol, we observed increased expression of *RDH10* (a key enzyme in embryonic retinol synthesis), *RBP4* (a hepatocyte-secreted retinol-binding protein), and *RARβ* (a nuclear receptor expressed in hepatic progenitors) between D3 and D20. This suggests that endogenous RA signaling was activated during hepatic development^45–47^ (Fig2I). Related biological terms were enriched in D20 samples (Supp. Fig. 2B). The expression of RA-responsive hepatic transcription factors *FOXA1* and *HNF1α*, which harbor conserved RA response elements, in both A10 and A100 aggregates (Supp. Fig. 2C) further supports this conclusion.

Regulatory network analysis of key pathway-associated genes and their targets revealed two major interaction clusters: one centered on hepatobiliary development (Cluster 1) and the other on organogenesis regulation (Cluster 2) (Fig. 2J). Gene ontology and tissue enrichment analyses confirmed strong associations with biological processes related to endoderm, mesenchyme, and hepatobiliary/cardiac tissue development (Fig. 2K, Supp Fig 2D).

Altogether, these findings indicate that our 3D differentiation platform supports the coordinated emergence of multiple cellular lineages, consistent with processes involved in organogenesis.

### Hepatic Cell Emergence Driven by Early Activin-Induced Differentiation

By day 20 of differentiation, aggregates derived from the DF6 hPSC line exhibited a transcriptional signature indicative of hepatic lineage commitment. Specifically, key hepatoblast markers such as *CK19* and *HNF4A* were expressed, confirming hepatic specification under both culture conditions (Fig. 3A).

**Figure 3.**
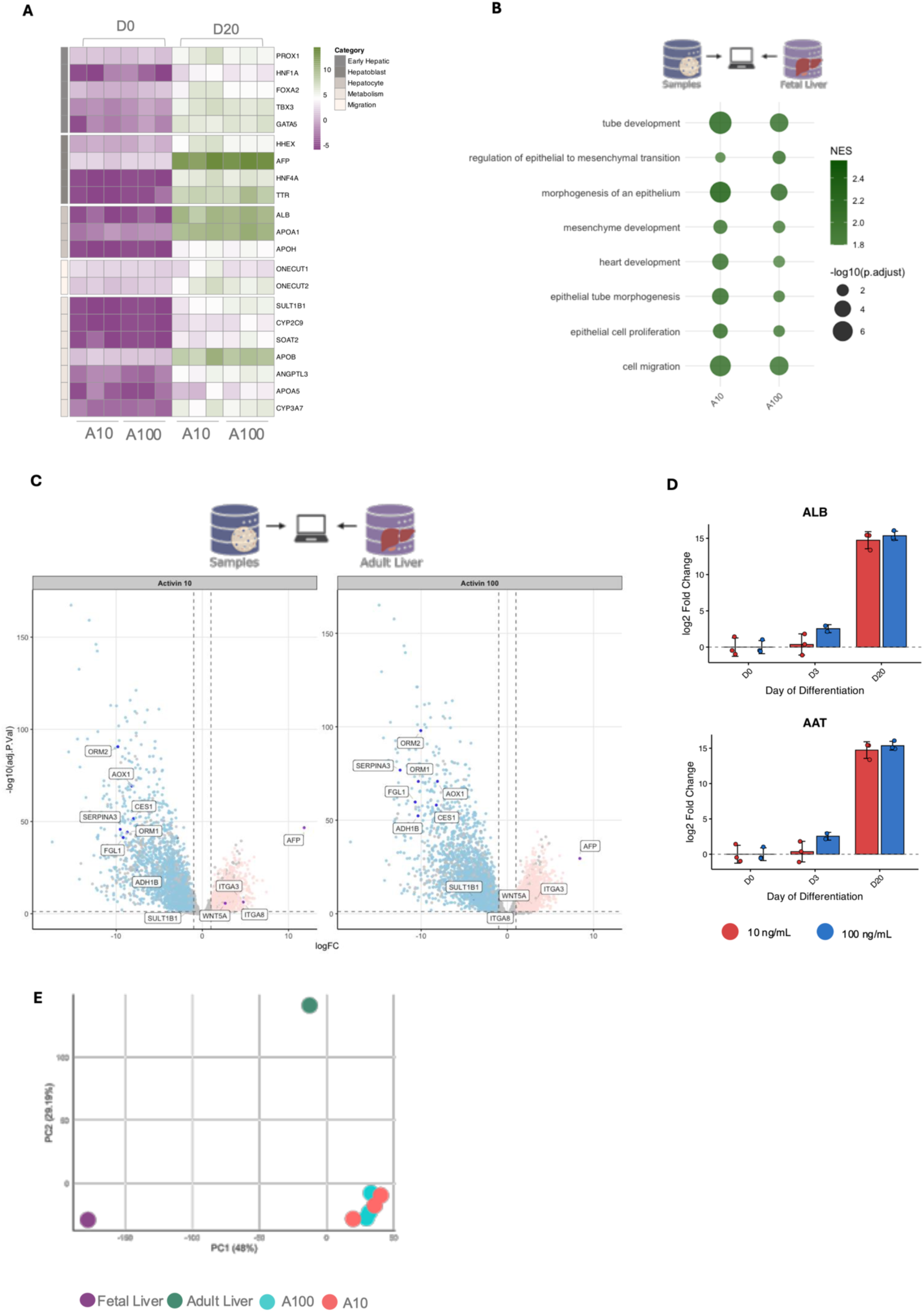
Molecular Profiling of early activin-induced differentiation revealed by Fetal/Adult molecular signatures comparisons. **(A) Gene expression heatmap during hepatic commitment:** Heatmap of hepatic commitment–associated genes showing log₂ counts per million (log₂CPM). Genes are organized by biological process or associated cell type, and columns represent sample conditions. Color intensity reflects expression level (low: dark green/teal; intermediate: light; high: purple/magenta). (B) **Gene set enrichment analysis relative to fetal liver:** Dot plot of enriched biological pathways in experimental samples compared to fetal liver. **(C) Differential gene expression in liver organoids under A10 and A100 conditions:** Volcano plots show –log₁₀ P versus log₂ fold change (FC) for all detected genes relative to adult liver. Left: A10 vs adult; right: A100 vs adult. Each point represents a gene; labeled genes indicate key liver maturation (*ORM2*, *AOX1*, *CES1*, *SERPINA3*, *ORM1*, *FGL1*, *ADH1B*, *SULT1B1*), mesodermal development (*WNT5A*, *ITGA3*, *ITGA8*) and fetal liver (*AFP*) markers. **(D) Expression dynamics of mature liver markers during hepatic differentiation:** Bar plots show mean ± S.D log₂ fold change (log₂ FC) in the expression of the mature hepatocyte markers **ALB** and **AAT** relative to day 0 (D0) for A10 (red) and A100 (blue) differentiation conditions (n=3) **(E) Principal component analysis of liver gene expression.** PCA of normalized gene expression shows PC1 (48% variance) versus PC2 (29% variance) across developmental stages and organoid conditions.

To assess the developmental fidelity of these organoids, we compared the transcriptomes of day 20 aggregates with publicly available human fetal and adult liver datasets. A heatmap comparison against fetal and adult liver references enabled the identification of the most relevant gene expression clusters for downstream analysis (Supp. Fig. 3A). Gene Set Enrichment Analysis (GSEA) of the most upregulated genes (relative to fetal liver) in both A10 and A100 samples revealed enrichment of biological processes associated with cell migration, epithelial morphogenesis, and tube formation (Fig. 3B). These features are consistent with the dynamic events of early liver development.

Because the reference fetal liver dataset represents embryos between 10 and 18 weeks of gestation, our D20 organoids likely correspond to an earlier developmental stage, approximately the third week of human gestation. At this stage, the ventral foregut epithelium thickens to form the liver diverticulum, which transitions from a monolayer of cuboidal endodermal cells to a multilayer of pseudostratified hepatoblasts that invade the septum transversum to form the liver bud^17^. Consistent with this, our organoids were enriched for mesoderm-associated biological processes, including cardiac development and epithelial transitions indicative of hepatoblast emergence (Fig. 3B).

Comparison with adult liver transcriptomes further confirmed the immature phenotype of both organoid conditions. Genes downregulated in A10 and A100 aggregates were primarily associated with mature liver functions such as lipid metabolism (*ORM2*, *FGL1, LGR1*), xenobiotic and alcohol metabolism (*AOX1*, *CES1*, *ADH1B*), and acute phase protein secretion (*SERPINA3*, *ORM1*) (Fig. 3C). In contrast, upregulated genes were enriched for developmental programs linked to mesoderm (*WNT5A, ITGA8, ITGA3*) differentiation. Notably, *AFP*, a well-established fetal hepatocyte marker^17^ was among the most highly upregulated genes in both conditions relative to adult liver, further reinforcing the fetal-like transcriptional profile.

Direct comparison between the two Activin A concentrations revealed dose-dependent differences in hepatic maturation. Expression levels of *ALB* and *AAT*, both markers of hepatic function, were significantly higher in A100 aggregates, while in A10 aggregates these markers were nearly undetectable (Fig. 3D). These findings suggest that higher Activin A levels promote greater hepatic functional maturation.

Conversely, the A10 condition was enriched for genes associated with hepatoblast migration and early hepatic or endodermal specification, including *ONECUT1/2*, *PROX1*, *HNF1A/B*, *FOXA2*, *HHEX*, *TBX3*, and *GATA5* (Fig. 3A). This gene expression profile indicates that A10 aggregates are developmentally delayed or maintained at a more primitive stage. In contrast, genes associated with hepatic metabolic function, such as *SULT1B1*, *CYP3A7*, *CYP2C9*, *APOA5*, and *ANGPTL3*, were significantly upregulated in A100 samples (Fig. 3A; Supp. Fig. 3B), supporting a more advanced hepatic phenotype in this condition.

Taken together, these results confirm that initial activin A induction effectively promotes endodermal specification, priming cells for hepatic differentiation. By day 20, the resulting hepatic-like cells under both conditions maintained an overall immature phenotype, most closely resembling embryonic or fetal liver tissue. Principal component analysis further supported this interpretation: the upper end of PC2, where adult reference was positioned, was enriched for genes associated with lipid, steroid, and drug metabolism, key features of hepatic maturation, whereas both our samples and fetal liver clustered at the lower end of PC2, consistent with an immature phenotype. (Fig3E, Supp Fig 3C). Pearson correlation analyses further corroborated the transcriptional proximity of both conditions to embryonic liver profiles (Supp. Fig. 3D).

Despite this overarching embryonic identity, aggregates cultured with 100 ng/mL Activin A demonstrated higher expression of hepatic metabolic genes (*CYP3A7*, *CYP2C9*, *ALB*) and sustained *HNF4A* levels, suggesting enhanced hepatic functional maturation. In contrast, aggregates exposed to 10 ng/mL Activin A maintained higher expression of genes related to progenitor migration and early hepatic/endodermal specification, such as *ONECUT1/2*, *PROX1*, *FOXA2*, and *HNF1A/B*, indicating a developmental profile consistent with earlier liver bud formation stages.

### Initial Low Activin A Concentrations Facilitate Liver Morphogenesis by Promoting the Emergence of Posterior Foregut-like Structures

To determine whether additional cell types beyond hepatic parenchyma emerged in A10 and A100 organoids, we analyzed RNA-seq data for activation of biological processes associated with mesodermal lineages, which are closely involved in liver development. To further assess cell type diversity, we performed gene deconvolution on the bulk RNA-seq profiles. This analysis revealed that both A10 and A100 aggregates contained hepatic parenchymal cells (hepatoblasts and hepatocytes), as well as non-parenchymal cells, including mesothelial and hematopoietic populations (Fig. 4A).

**Figure 4.**
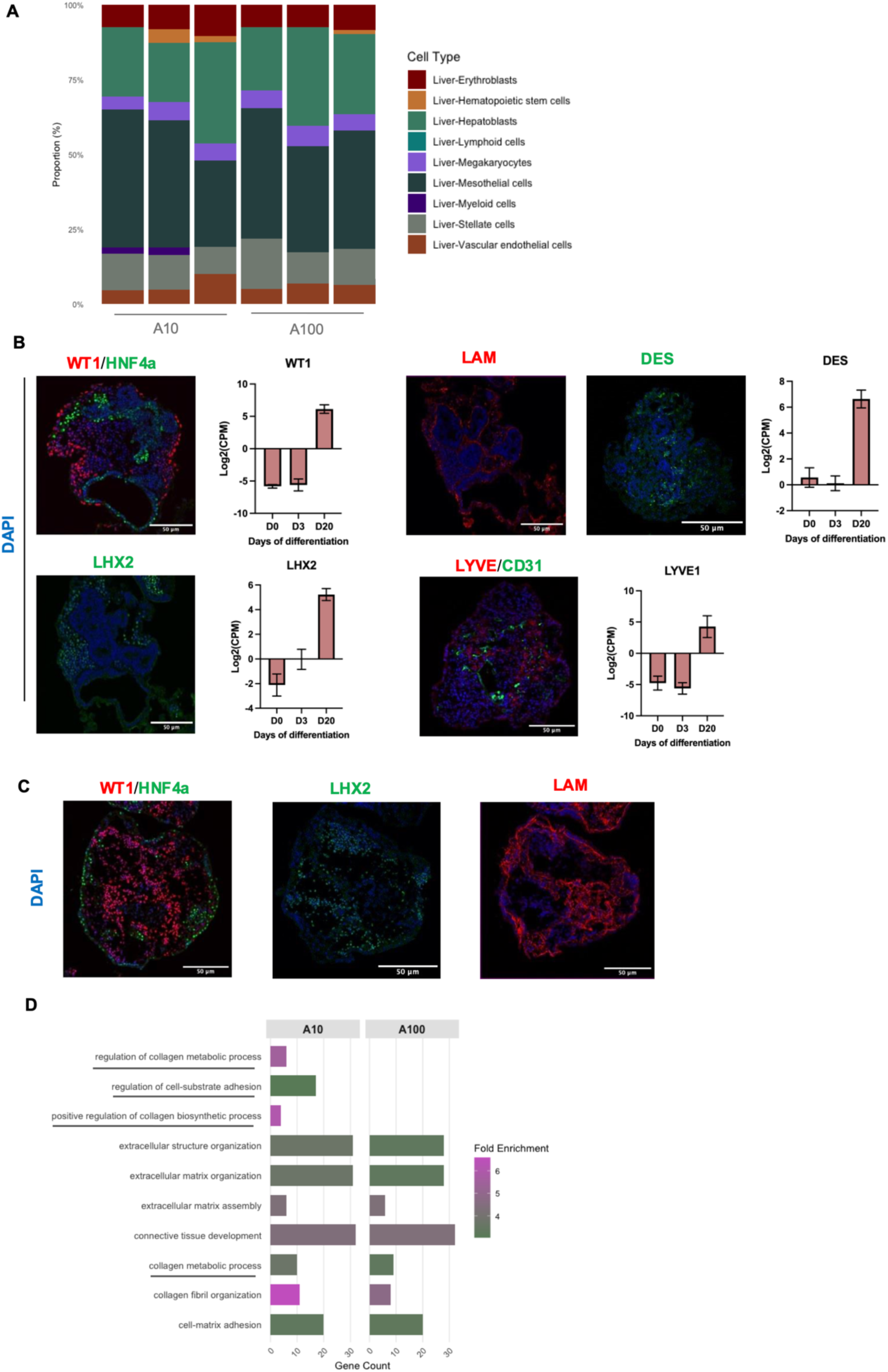
Integrative Cellular and Molecular profiling reveals activin-dependent foregut-like composition and ECM remodeling. **(A) Cell-type composition of bulk RNA-seq samples inferred by deconvolution:** Stacked bar plots show proportions of parenchymal and non-parenchymal cell populations inferred from bulk RNA-seq of liver organoids cultured under A10 and A100 conditions (n=3) using the DeconRNASeq package. Colors denote distinct cell types. **(B) Immunofluorescence and gene expression analysis of liver organoids cultured under A10 condition:** Representative images show staining for hepatocytes (HNF4A, green), hepatic mesothelium (WT1, red; DES, green), hepatic endothelium (LYVE1, red; CD31, green), laminin (LAM, red) and hepatic mesenchymal cells (LHX2, green). Nuclei are counterstained with DAPI (blue). Scale bars 50 𝝁m. Corresponding bar plots show mean log₂(CPM) expression of the same markers from RNA-seq (n = 3), with error bars indicating SD. **(C) Immunofluorescence analysis of liver organoids cultured under A100 condition:** Confocal images show hepatic mesothelium (WT1, red), hepatocytes (HNF4A, green), mesenchymal cells (LHX1, green), and basement membranes (LAM, red). Scale bars 50 𝝁m **(D) GO enrichment of liver organoids in A10 vs. A100 conditions:** Bar plot of significantly enriched Biological Process GO terms (Fold Enrichment >1, p < 0.05) related to collagen metabolism and ECM. The y-axis shows GO terms, the x-axis shows gene count, and bar color represents Fold Enrichment, highlighting differences between the two culture conditions.

Brightfield imaging revealed distinct tube-like structures in A10 aggregates, which were less frequently observed in A100 samples (Fig. 1C). Given that both conditions appeared to generate similar cell types, we hypothesized that the primary difference between A10 and A100 organoids lies in their cellular organization rather than composition.

Day 20 cryosections confirmed significant differences in spatial organization at later stages of differentiation. A10 aggregates displayed lumen-like structures reminiscent of the early steps of liver bud morphogenesis seen during embryogenesis which persisted throughout extended culture (Fig. 4B, Supp Fig 4A). HNF4α⁺ cells were arranged into epithelial lumens resembling posterior foregut-like tubes. These lumens were separated by a LAM⁺ (laminin-positive) basal lamina from adjacent LHX2⁺ cells, analogous to hepatic mesenchymal cells (HMCs) surrounding the liver bud *in vivo*. Additionally, WT1⁺/DES⁺ cells were observed at the periphery of the organoids, alongside CD31^+^/LYVE1⁺ cells, potentially representing nascent hepatic mesothelium and endothelial structures (Fig. 4B). RNAseq data showed significantly higher expression of these markers at D20 in A10 conditions (Fig. 4B) and the trend across all differentiation timepoints was confirmed by qRT-PCR (Supp Fig 4B), sustaining the enhanced emergence of these supporting mesenchymal lineages.

In contrast, although non-parenchymal cells were also detected in A100 aggregates, they were less abundant and lacked the spatial and structural organization observed in A10 (Fig. 4C). Basal lamina structures were more prominent in A10 aggregates, consistent with a higher enrichment of biological processes related to collagen metabolism compared to A100 (Fig. 4D). This aligns with previous studies indicating that fetal liver cell types synthesize key extracellular matrix components, such as collagens I, III, IV, and laminin^48,49^.

Together, these findings indicate that while both A10 and A100 conditions generate parenchymal and non-parenchymal lineages, lower Activin A concentrations promote enhanced spatial organization of non-parenchymal compartments. This environment is more permissive for recapitulating the coordinated morphogenetic events characteristic of liver bud formation during early embryonic development.

## Discussion

### Recapitulating Morphogen Gradients *In Vitro*: Coordinating Endoderm–Mesoderm specification in Liver Organoids

*In vivo* liver development requires the integration of spatially and temporally regulated signals between the definitive endoderm (DE) and adjacent mesoderm. During early embryogenesis, a gradient of Nodal signaling directs anterior endoderm formation, with higher Nodal levels promoting *FOXA2* expression and commitment to anterior definitive endoderm (ADE). DE progenitors migrate to juxtapose the cardiac mesoderm and septum transversum mesenchyme (STM), which secrete instructive factors, such as FGFs, WNTs, BMP4, and retinoic acid, in a concentration-dependent manner to refine endodermal regional identity^16,17^ . Following specification, the hepatic endoderm thickens into a pseudostratified diverticulum. Hepatoblasts then delaminate, proliferate, and invade the STM, processes critically dependent on coordinated epithelial–mesenchymal crosstalk and extracellular matrix remodeling ^50–52^.

John Gurdon’s seminal work demonstrated that cells activate specific gene expression programs based on their distance from a localized morphogen source, establishing the foundation for gradient-based developmental patterning^23^. Despite the development of various *in vitro* strategies, replicating the precise timing and coordination of these morphogenic interactions remains a significant challenge. Three-dimensional differentiation systems using hPSCs have proven more effective at recapitulating aspects of embryogenesis, particularly by enabling the formation of morphogen gradients^53,54^.

Building on this principle, we designed a 3D co-emergence platform that supports the simultaneous generation of DE and STM lineages from hPSCs. By modulating Activin A concentrations, specifically using 10 ng/mL (A10) to favor mesendodermal differentiation, and 100 ng/mL (A100) to drive uniform endoderm specification, we aimed to mimic *in vivo* Nodal/Activin gradients and investigate their influence on cell fate decisions and spatial patterning.

Consistent with the gradient theory, A10 aggregates exhibited peripheral enrichment of SOX17 and TBX3, with diminishing expression toward the center, while A100 aggregates showed a more homogeneous distribution of these markers. This pattern parallels *in vivo* Nodal dynamics, where lower Nodal levels favor mesoderm specification and higher levels support definitive endoderm induction^55^. Temporal gene expression analysis further confirmed this trajectory: early upregulation of *SOX17* and *TBX3* was followed by the emergence of hepatoblast and hepatocyte markers, including *HNF4A* and *AFP*, aligning with established timelines of liver development^2,9,30^ (Fig. 5).

**Figure 5.**
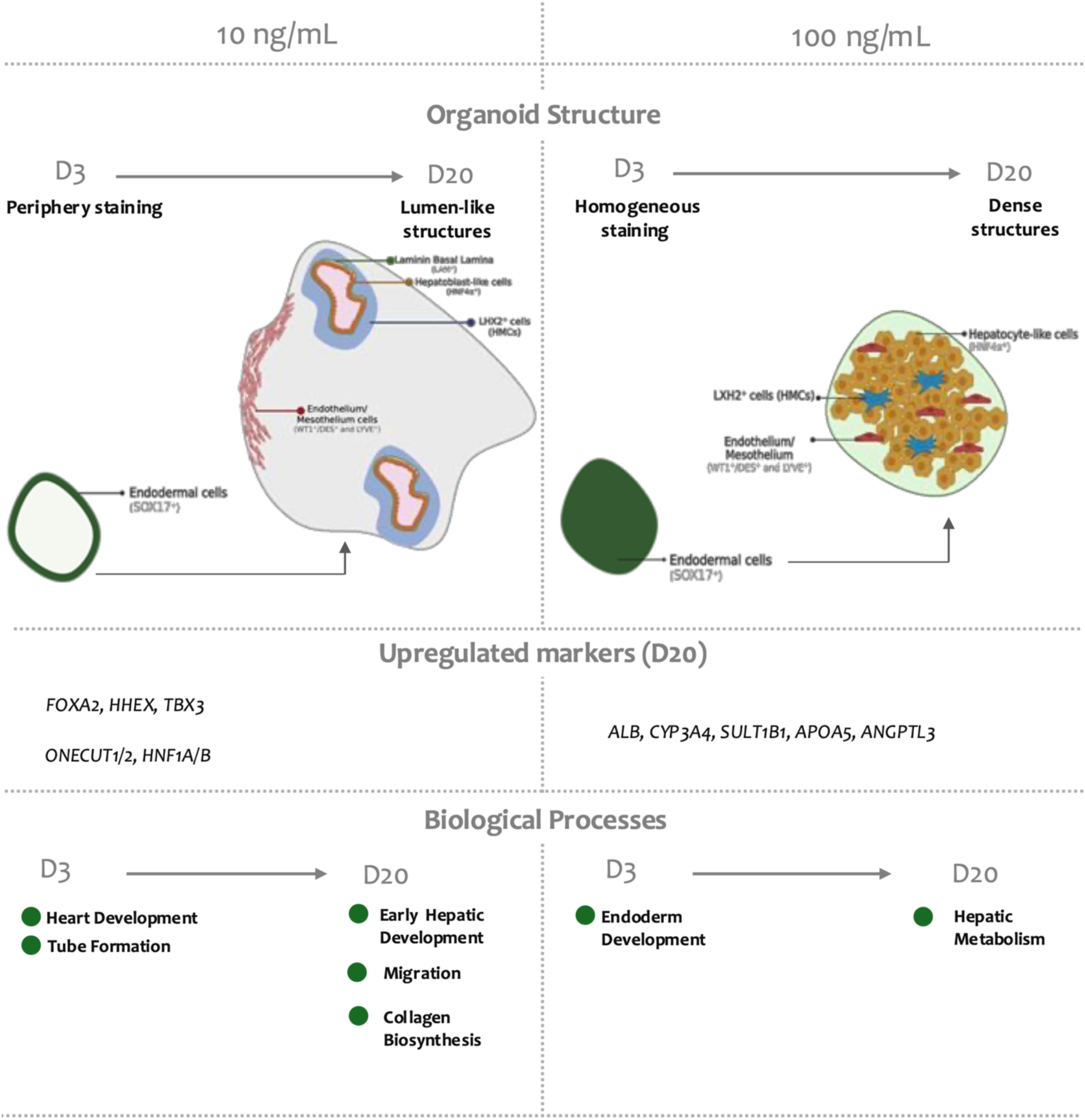
Organoid development under different growth factor concentrations: Organoids cultured with 10 ng/mL (left) or 100 ng/mL (right) Activin A show distinct structural and molecular trajectories from day 3 (D3) to day 20 (D20). At 10 ng/mL, peripheral staining at D3 evolves into lumen-like structures with hepatoblast-like cells, endothelium, and mesenchymal cells. At 100 ng/mL, homogeneous staining at D3 transitions into dense structures enriched in hepatocyte-like cells. Upregulated markers at D20 include *FOXA2*, *HHEX*, *TBX3*, *ONECUT1/2*, and *HNF1A/B* at 10 ng/mL, and *ALB*, *CYP3A4*, *SULT1B1*, *APOA5*, and *ANGPTL3* at 100 ng/mL. Associated biological processes shift from early hepatic development, migration, and collagen biosynthesis at 10 ng/mL to hepatic metabolism at 100 ng/mL.

### Co-Emergence of Mesenchymal Lineages and Hepatic Niche Formation

Cell–cell communication within the forming organoid is essential, as failure to establish cooperative and synergistic interactions can disrupt tissue development^34^. Several published protocols have demonstrated that hPSC-derived progenitors can generate distinct embryonic germ layers within a single organoid^56–59^.

Here, we observed upregulation of TGF-β/Nodal/Activin signaling at day 3 and sustained WNT activity through day 20, consistent with the emergence of mesenchymal lineages. The presence of LHX2⁺ cells surrounding tubular structures and WT1⁺ cells in A10 aggregates supports the autonomous generation of non-parenchymal mesenchymal populations. Expression of *FGF8*, *FGF10*, along with BMP ligands and downstream targets *GATA4/6* at day 3, mirrors early mesodermal instructive signaling. Notably, *FGF10* remained expressed through day 20, consistent with its established role in hepatoblast proliferation and survival^43,44^.

BMP signaling, together with other STM-derived mesenchymal signals, is critical for hepatic induction. Inhibition of BMP can shift the anterior hepato-pancreatic boundary toward a pancreatic fate^43,60^. These findings further support the idea that our co-emergence system enables the generation of mesodermal lineages involved in liver development. Additionally, both A10 and A100 conditions showed not only activation of these signaling pathways, but also their regulation, as evidenced by the expression of pathway inhibitors and enrichment of relevant biological processes.

Morphological divergence between aggregates was also observed. A10 cultures developed cystic, bud-like structures reminiscent of liver diverticula, whereas A100 aggregates remained dense and reticulated. This phenotypic difference reinforces the importance of gradient-dependent mesendodermal interactions for tissue morphogenesis and liver bud emergence, similar to co-culture organoid systems that rely on stromal FGF/BMP signaling^30^. Interestingly, the bud-like spatial organization seen in A10 samples resembles the morphologies observed in hydrogel-based liver organoids, where internal lumen formation is commonly detected^61^. In our system, the appearance of a basal lamina surrounding epithelial buds suggests the establishment of hepatocyte polarity, a critical feature for liver-specific function, akin to what has been reported in hydrogel culture systems^62^.

Hydrogels are known to recapitulate key biomechanical and biochemical cues that guide cellular self-organization and differentiation. They can effectively influence cell fate decisions and support complex tissue architecture^61,62^. While marker-based confirmation is still required to definitively demonstrate polarity in our model, the observed structural features are compelling. Notably, our co-emergence system appears to achieve comparable morphogenetic outcomes without the extensive and labor-intensive processes typically associated with hydrogel use, suggesting a simpler yet effective alternative for hepatic tissue engineering.

The retinoic acid (RA) pathway represents another key mesenchymal signaling axis during liver development. In mice, RA synthesis begins at embryonic day 7.5 with *RDH10* expression in the presomitic mesoderm and plays a crucial role in terminal hepatic differentiation. RA inhibition at pre-hepatic stages (E8.25) leads to loss of the anterior liver bud, while RA excess disrupts the entire liver bud, underscoring the importance of precisely timed and spatially regulated RA activity^45–47,63^. Branco et al. showed that RA pathway enhancement influences organoid development: WNT and BMP alone promote early hepatic specification, while early RA supplementation supports multilineage organization, including pro-epicardium, STM, and posterior foregut^57^. In our system, we observed increasing expression of *RDH10*, *RBP4*, and *RARβ*, indicating a gradual, temporally controlled, and autonomous activation of the RA pathway, without exogenous supplementation, consistent with endogenous developmental mechanisms^47^.

Finally, regulatory network analysis revealed two major clusters: one associated with hepatobiliary development and another related to general organogenesis. Gene ontology and tissue-specific enrichment analyses further aligned these networks with endodermal and mesenchymal tissues, as well as hepatobiliary and cardiac development. Collectively, these findings suggest that our model recapitulates aspects of the signaling interactions involved in endoderm regionalization and hepatic niche formation.

### Activin-Induced Organoids Recapitulate Early Liver Bud Formation and Fetal Hepatic Traits

Our bulk RNA-seq data further confirmed that initial induction with Activin A effectively promoted both endodermal and hepatic differentiation. At day 20 (D20), organoids from both conditions exhibited transcriptomic profiles more closely resembling embryonic liver tissue. This was supported by the downregulation of mature liver function genes, specifically, those involved in lipid metabolism (*e.g.*, *ORM2, FGL1*), drug and alcohol metabolism (*e.g.*, *AOX1, CES1, ADH1B*), and acute-phase protein secretion (*e.g.*, *SERPINA3, ORM2*). Concurrently, there was an upregulation of developmental and hepatic fetal markers, such as *AFP,* supporting the immature hepatic phenotype of the organoids.

Hierarchical clustering revealed enrichment for mesoderm-related biological processes, including genes associated with heart development, cell migration, and epithelial transitions (*e.g.*, from cuboidal to pseudostratified states). These transcriptional signatures suggest that our organoids share molecular features with early stages of hepatic specification. This molecular profile is consistent with early developmental stages when the ventral foregut thickens to form the liver diverticulum. During this period, endodermal cells transition into proliferative hepatoblasts that invade the septum transversum, initiating liver bud formation^16,17^. These findings are consistent with the observations of Takebe and colleagues in their study on co-cultured liver bud organoids, where the day 4 (D4) transcriptomic profile, based on 83 serially upregulated liver developmental genes, closely matched that of mouse liver buds at embryonic days E10.5–E11.5 (approximately equivalent to the third or fourth week of human gestation), rather than more advanced fetal or adult liver tissue^30^.

Interestingly, when comparing A10 and A100 samples, we observed differences in maturation states. A100 aggregates exhibited significantly higher expression of mature hepatic markers, including AAT and ALB, as well as metabolic genes such as *SULT1B1*, *CYP3A7*, *CYP2C9*, *APOA5*, and *ANGPTL3*. In contrast, A10 samples showed enrichment for genes involved in hepatoblast migration and early hepatic/endodermal specification.

It is important to acknowledge the limitations of our study. We did not perform single-cell transcriptomics, which would provide insights into cellular heterogeneity, lineage composition, and maturation states of both parenchymal and non-parenchymal populations. Functional hepatic assays, such as albumin secretion or cytochrome P450 activity, were also not conducted, so the physiological functionality of the differentiated cells remains to be confirmed. Additionally, we did not assess long-term culture or maturation, leaving open whether the model can support later stages of liver development or maintain hepatic identity over extended periods.

### Harnessing Morphogen Gradients for Cross-Lineage Organoid Specification

Our findings also underscore how early morphogen gradients can be harnessed to generate ventral foregut organoids beyond the liver, particularly pancreas, gallbladder, and thyroid, by fine-tuning Activin A to guide mesendodermal patterning^64^. The development and optimization of liver organoid cultures provide a valuable framework for modeling other gut tube–derived organs, which similarly arise as epithelial buds surrounded by mesodermal derivatives. Paracrine BMP, FGF, and WNT signals from adjacent mesenchymal populations at critical developmental windows (*e.g.*, E8.0 in mouse) help distinguish organ domains^43^ offering a blueprint for engineering other organ systems *in vitro*.

## Conclusion

In this study, we demonstrate that low versus high Activin A concentrations in a 3D co-emergence platform recreate essential Nodal/Activin gradients, enabling autonomous generation of hepatic and mesenchymal lineages, recapitulating key stages of liver bud development. Our findings suggest the potential applicability of morphogen-guided differentiation strategies for generating ventral foregut-derived organoids. By leveraging fine-tuned gradients of Activin A and coordinating mesendodermal patterning, it is possible to direct hPSCs toward multiple gut tube lineages beyond the liver. Thus, this gradient-based strategy provides a versatile framework for modeling endoderm–mesoderm interactions across multiple gut-tube derivatives (*e.g.*, pancreas, gallbladder, or thyroid), advancing both developmental biology and organoid engineering.

## Author Contributions

All authors contributed to the conceptualization and experimental design of the study. JPC performed all cell culture experiments and imaging acquisition. ACB carried out all data processing, quantitative analyses, and image analysis. JPC, ACB, and TGF jointly interpreted the results and wrote the manuscript. DESN, CAVR, and JMSC provided critical feedback throughout the project and contributed to manuscript revision. All authors reviewed and approved the final version of the manuscript.

## Acknowledgements

We thank Gulbenkian Institute for Molecular Medicine (GIMM) for the access to the histopathology and bioimaging facilities for organoid sample processing and imaging. We acknowledge Nuno Morais Lab for providing the processed adult liver data obtained from their voyAGEr platform. We acknowledge PBS Biotech, Inc. for providing the PBS Mini vertical-wheel bioreactors used in this study. We acknowledge funding received from Fundação para a Ciência e a Tecnologia (FCT), through Institute for Bioengineering and Biosciences (projects UIDB/04565/2020 and UIDP/04565/2020) and through Associate Laboratory Institute for Health and Bioeconomy (LA/ P/0140/2020). FCT is also acknowledged for the PhD grants to JPC and ACB (SFRH/PD/BD135500/2018 and UI/BD/153364/2022, respectively).

## Supplementary Figures

**Supplementary Fig 1A.**
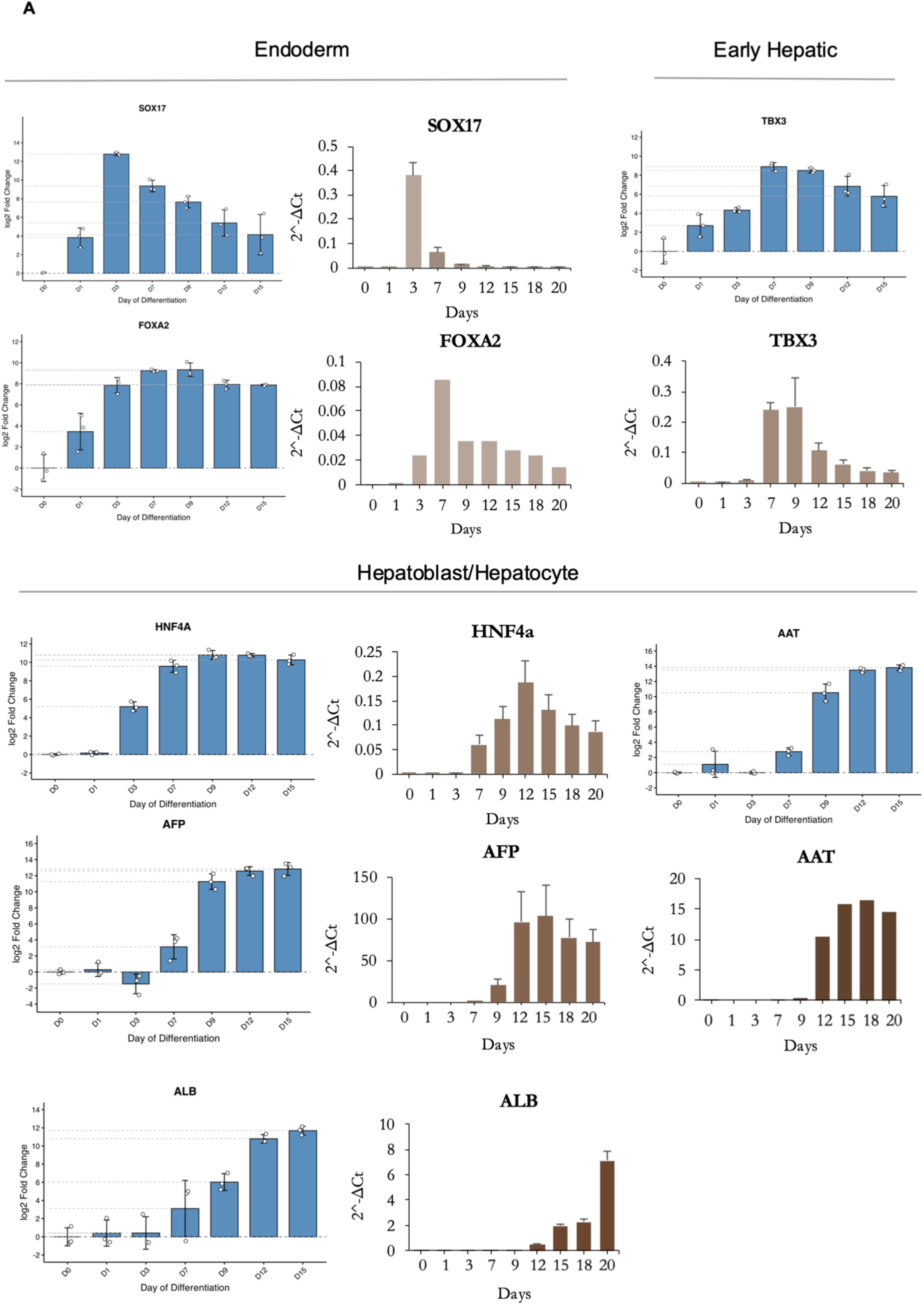
Validation of lineage marker expression during 2D hepatic differentiation. RNA-seq expression profiles were validated by quantitative PCR (qPCR) for representative lineage markers across differentiation stages (day 0–20). Expression of definitive endoderm markers (*SOX17*, *FOXA2*), early hepatic marker (*TBX3*), and hepatoblast/hepatocyte markers (*HNF4A*, *AFP*, *ALB*, *AAT*) was assessed. Blue bars indicate RNA-seq expression (log₂ fold change ± error, *n* = 3), and brown bars indicate qPCR expression (2^–ΔCt ± error).

**Supplementary Fig 1B.**
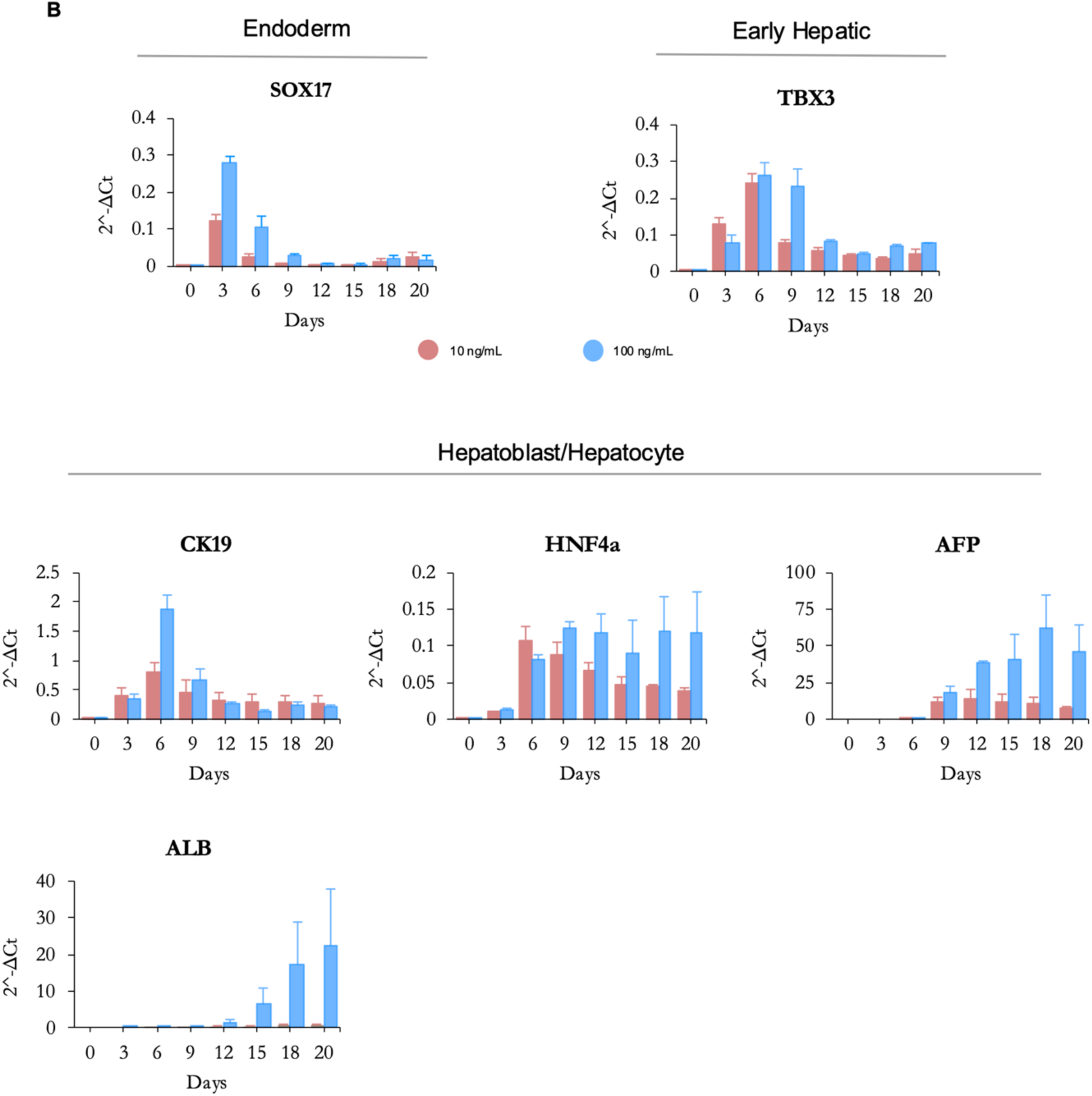
qPCR analysis of lineage marker expression during 3D hepatic differentiation: Expression of endoderm (*SOX17*, *FOXA2*), early hepatic (*TBX3*), and hepatoblast/hepatocyte (*HNF4A*, *AFP*, *ALB*, *CK19*) markers was assessed from day 0 to day 20. Bars represent 2^–ΔCt values ± error.

**Supplementary Fig 2.**
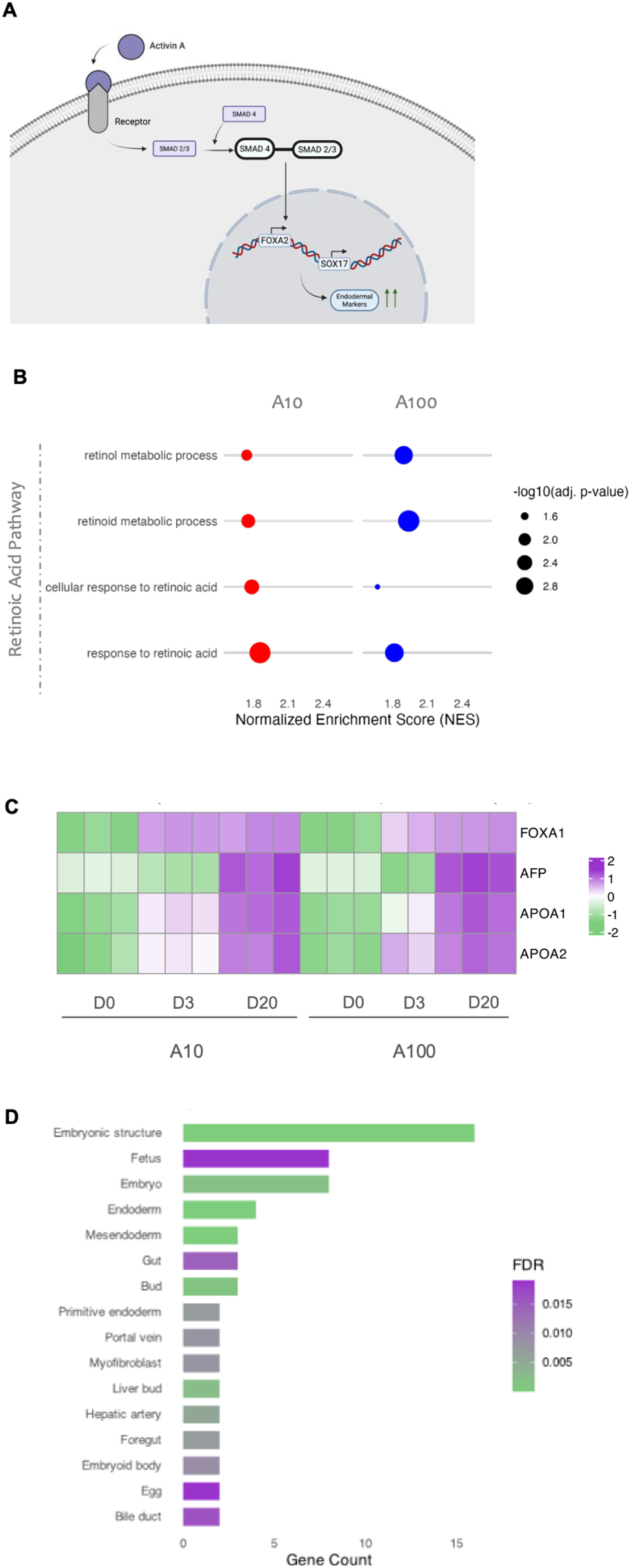
(A) Activin A–induced Nodal signaling and endoderm specification: Diagram of Activin A–induced Nodal signaling driving SOX17/FOXA2 activation and endodermal fate specification. **(B) GSEA of Retinoic Acid pathway–related GO terms between A10 and A100:** Dot plot of terms passing multiple-testing correction (adjusted *p* < 0.05). NES is plotted on the x-axis, GO terms on the y-axis, and dot size represents –log₁₀(adjusted *p* value). **(C) Retinoic acid–responsive gene expression during organoid differentiation:** Heatmap of log₂ expression levels of retinoic acid–responsive genes (*FOXA1*, *AFP*, *APOA1*, *APOA2*) across differentiation timepoints (D0, D3, D20) in A10 and A100 conditions. **(D) Tissue enrichment of network-derived genes:** Tissue enrichment analysis of differentially expressed genes from the protein–protein interaction network. Bars indicate significantly enriched tissues (FDR < 0.05), with length corresponding to gene count.

**Supplementary Figure 3.**
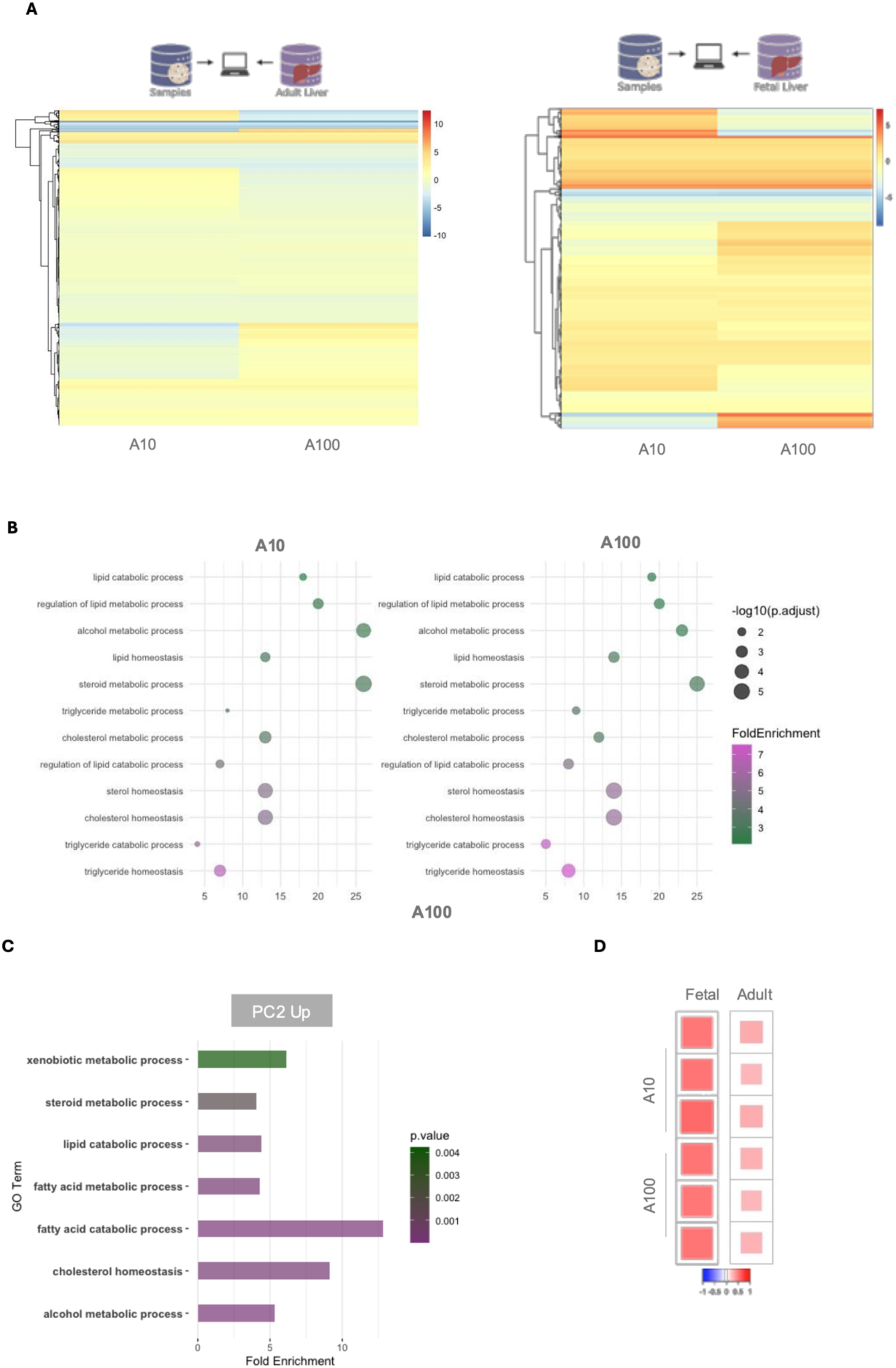
(A) Differential gene expression in D20 liver organoids relative to fetal and adult liver. Heatmap of log₂ fold change for genes significantly altered (adjusted P < 0.05) in A10 and A100 organoids at D20. Rows: hierarchically clustered genes; columns: biological replicates (n = 3). Red: upregulated; blue: downregulated. **(B) Gene Ontology enrichment analysis of hepatic metabolism:** Dot plot showing GO pathways significantly enriched in experimental samples. **(C) Gene Ontology enrichment of PC2 top-contributing genes.** Bar plot shows GO terms enriched in the adult dataset for genes driving PC2 (fold enrichment >1, P < 0.05). **(D) Transcriptional correlation between liver organoids and reference datasets.** Pearson correlation matrix of normalized gene expression profiles (n = 3 per condition) for A10 and A100 organoids versus fetal and adult liver. Color intensity reflects correlation strength (red = strong positive, white = none).

**Supplementary Fig 4.**
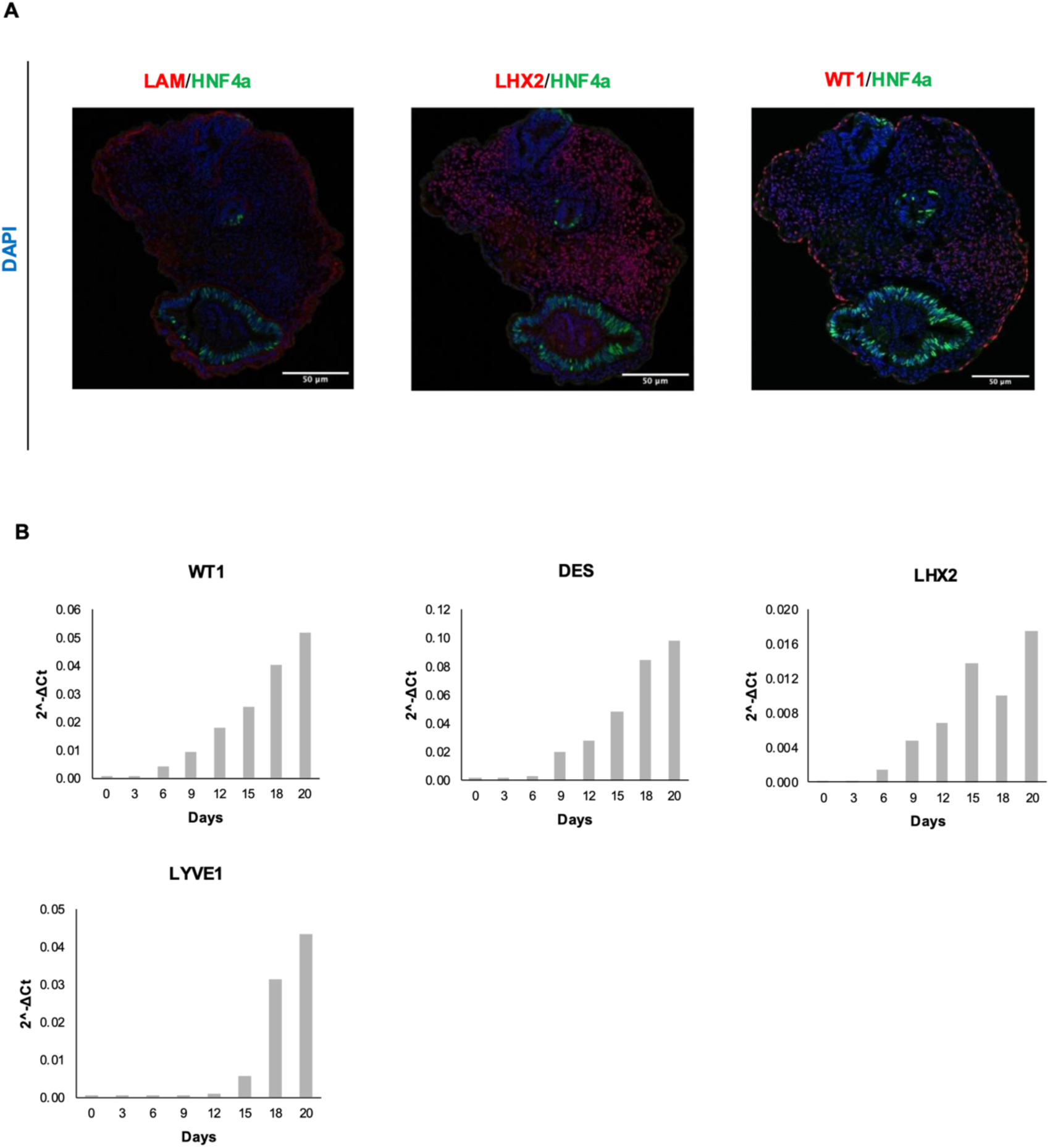
(A) Immunofluorescence analysis of D40 cryosectioned human liver organoids: Confocal images show lumen-like structures in human liver organoids. HNF4α (green) marks hepatocytes, while the red signal highlights different cell types in each panel: Laminin (LAM, left) outlines the basement membrane around the epithelial lumen; LHX2 (middle) marks hepatic mesenchymal cells outside the HNF4α-positive lumens; and WT1 (right) labels mesothelial cells surrounding the organoid structure. Nuclei are counterstained with DAPI (blue) to show overall cellular organization. Scale bars 50 𝝁m **(B) qPCR analysis of non-parenchymal liver markers’ expression *(WT1, DES, LHX2, LYVE1)* during 3D hepatic differentiation in A10 conditions:** Bars represent 2^–ΔCt values

**Supplementary Table 1:**
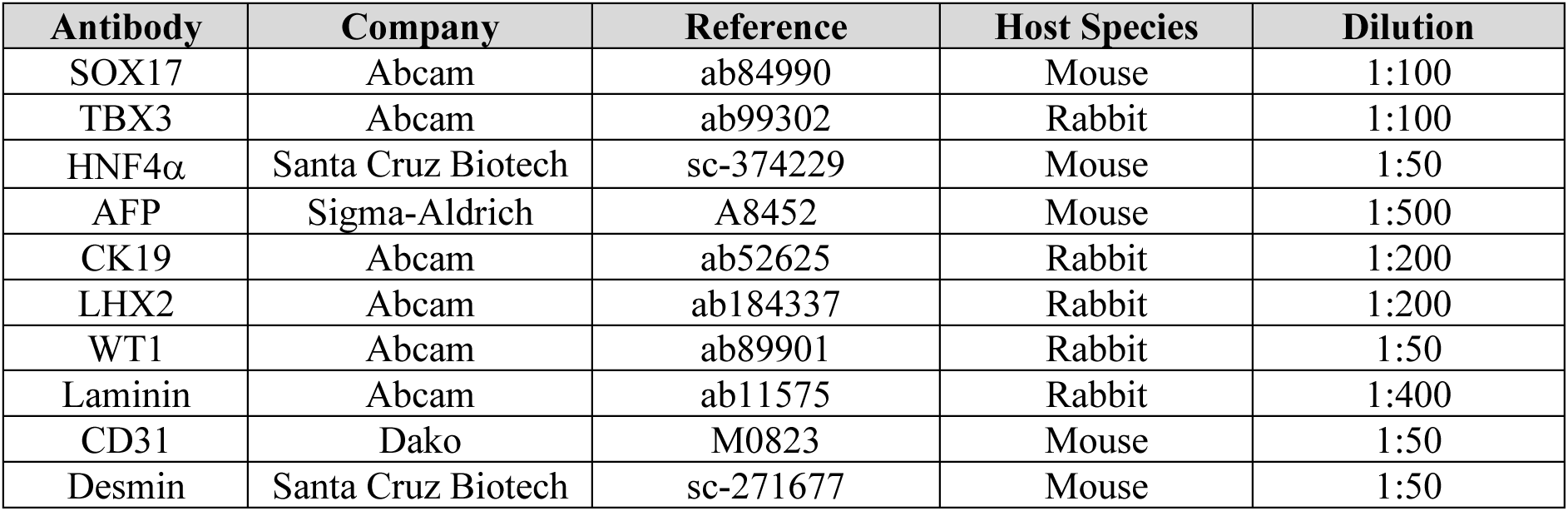
List of primary antibodies and dilutions used for immunostaining.

**Supplementary table 2:**
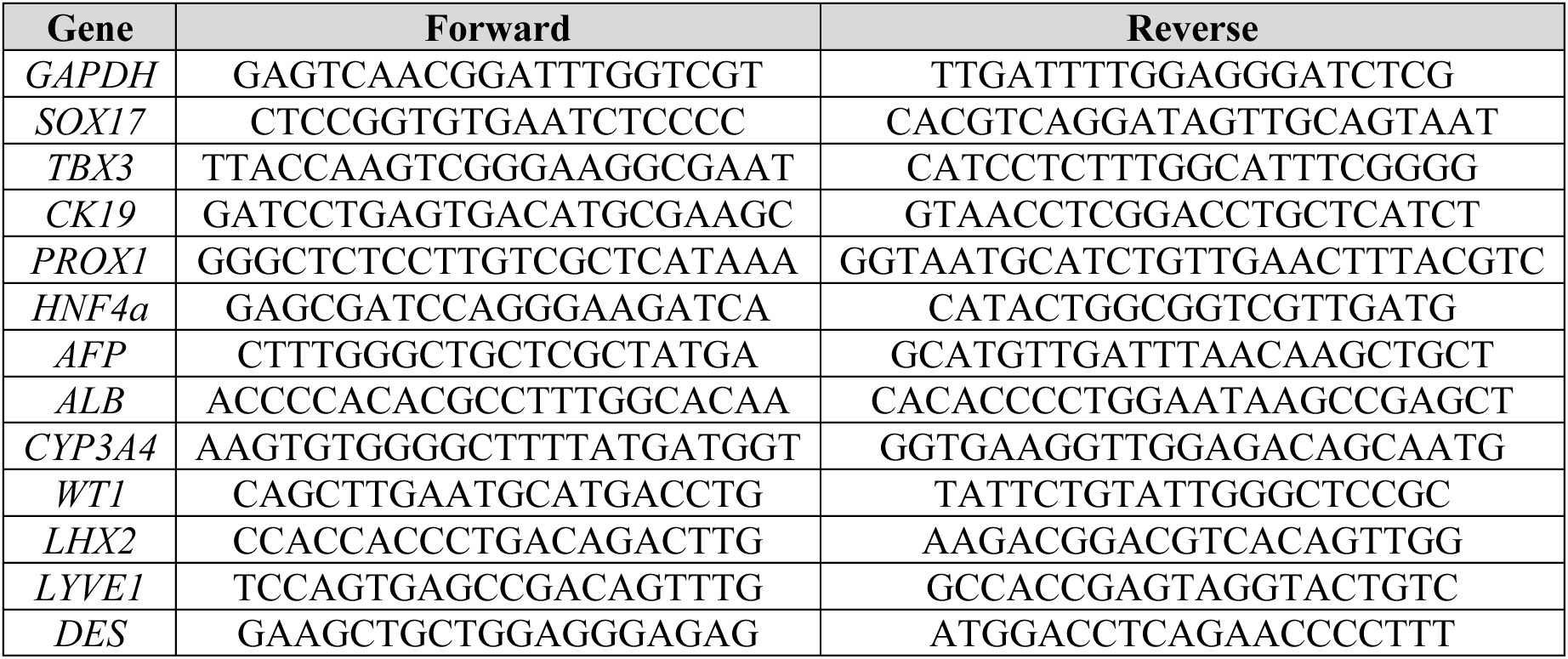
List of Primers.

