## Supplemental Tables for "Gradient-Based Regulation of Activin A Directs Multilineage Liver Organoid Development from Human Pluripotent Stem Cells"

Supplementary Table 1: List of primary antibodies and dilutions used for immunostaining

| Antibody | Company | Reference | Host Species | Dilution |
|---|---|---|---|---|
| SOX17 | Abcam | ab84990 | Mouse | 1:100 |
| TBX3 | Abcam | ab99302 | Rabbit | 1:100 |
| HNF4$\alpha$ | Santa Cruz Biotech | sc-374229 | Mouse | 1:50 |
| AFP | Sigma-Aldrich | A8452 | Mouse | 1:500 |
| CK19 | Abcam | ab52625 | Rabbit | 1:200 |
| LHX2 | Abcam | ab184337 | Rabbit | 1:200 |
| WT1 | Abcam | ab89901 | Rabbit | 1:50 |
| Laminin | Abcam | ab11575 | Rabbit | 1:400 |
| CD31 | Dako | M0823 | Mouse | 1:50 |
| Desmin | Santa Cruz Biotech | sc-271677 | Mouse | 1:50 |

Supplementary table 2: List of Primers

| Gene | Forward | Reverse |
|---|---|---|
| GAPDH | GAGTCAACGGATTTGGTCGT | TTGATTTTGGAGGGATCTCG |
| SOX17 | CTCCGGTGTGAATCTCCCC | CACGTCAGGATAGTTGCAGTAAT |
| TBX3 | TTACCAAGTCGGGAAGGCGAAT | CATCCTCTTTGGCATTTCGGGG |
| CK19 | GATCCTGAGTGACATGCGAAGC | GTAACCTCGGACCTGCTCATCT |
| PROX1 | GGGCTCTCCTTGTCGCTCATAAA | GGTAATGCATCTGTTGAACTTTACGTC |
| HNF4a | GAGCGATCCAGGGAAGATCA | CATACTGGCGGTCGTTGATG |
| AFP | CTTTGGGCTGCTCGCTATGA | GCATGTTGATTTAACAAGCTGCT |
| ALB | ACCCCACACGCCTTTGGCACAA | CACACCCCTGGAATAAGCCGAGCT |
| CYP3A4 | AAGTGTGGGGCTTTTATGATGGT | GGTGAAGGTTGGAGACAGCAATG |
| WT1 | CAGCTTGAATGCATGACCTG | TATTCTGTATTGGGCTCCGC |
| LHX2 | CCACCACCCTGACAGACTTG | AAGACGGACGTCACAGTTGG |
| LYVE1 | | |
| DES | | |
